# Eph/ephrin signalling in the developing brain is regulated by tissue stiffness

**DOI:** 10.1101/2024.02.15.580461

**Authors:** Jana Sipkova, Kristian Franze

## Abstract

Eph receptors and their membrane-bound ligands, ephrins, provide key signals in many biological processes, such as cell proliferation, cell motility and cell sorting at tissue boundaries. However, despite immense progress in our understanding of Eph/ephrin signalling, there are still discrepancies between *in vitro* and *in vivo* work, and the regulation of Eph/ephrin signalling remains incompletely understood. Since a major difference between *in vivo* and most *in vitro* experiments is the stiffness of the cellular environment, we here investigated the interplay between tissue mechanics and Eph/ephrin signalling using the *Xenopus laevis* optic pathway as a model system. *Xenopus* retinal neurons cultured on soft substrates mechanically resembling brain tissue showed the opposite response to ephrinB1 compared to those cultured on glass. *In vivo* atomic force microscopy (AFM)-based stiffness mapping revealed that the visual area of the *Xenopus* brain, the optic tectum, becomes mechanically heterogeneous during its innervation by axons of retinal neurons. The resulting stiffness gradient correlated with both a cell density gradient and expression patterns of EphB and ephrinB family members. Exposing *ex vivo* brains to stiffer matrices or locally stiffening the optic tectum *in vivo* led to an increase in EphB2 expression in the optic tectum, indicating that tissue mechanics is an important regulator of Eph/ephrin signalling. Similar mechanisms are likely to be involved in the development and diseases of many other organ systems.

---

Ephrins and Eph receptors are transmembrane proteins that interact with each other to enable contact-mediated interactions between cells. There are A and B subclasses of both proteins, depending on their sequences and binding affinities to their respective A– or B-type binding partners. Although Ephs and ephrins are classically referred to as receptor and ligand, respectively, upon binding, signalling can occur downstream of either binding partner (Fig. S1a). Additional complexity in the signalling pathway arises from interactions between family members within the same cell and cellular processes such as receptor clustering, proteolytic cleavage, endocytosis and local protein translation^1,2^. Eph/ephrin signalling is therefore highly versatile and involved in many physiological processes, from neural development to bone homeostasis, as well as in pathologies such as cancer^3^.

Due to the diversity of signalling modes and downstream pathways, the regulation of Eph/ephrin signalling is not yet fully understood. This is exemplified by studies of the retinotectal projection, where Ephs and ephrins in retinal ganglion cells (RGCs) and the visual area of the brain, the optic tectum, help to establish precise connections between the eye and the brain^4,5^ (see Fig. S1b, c for a schematic). This process, known as retinotectal mapping, has primarily been studied using *in vitro* explant cultures^1,6–11^ and *in vivo* mutagenesis^12–15^. The results of these studies are often conflicting, particularly in the case of EphB/ephrinB signalling, which is thought to be responsible for retinotectal mapping along the dorso-ventral (D-V) axis. In particular, while it has been consistently reported that axons of ventral and dorsal RGCs respond differentially to ephrinB cues *in vitro*^1,8,9^, disruption of signalling downstream of EphB *in vivo* does not affect the guidance of ventral or dorsal RGC axons to their initial positions in the optic tectum^12,13^. Furthermore, despite extensive research into the mechanisms that determine the attractive versus repulsive effects of Eph receptors and ligands, it remains unclear how these effects are regulated *in vivo*. For instance, ephrinB1 is repulsive *in vitro*^1,9^, but *in vivo* it serves as both an attractive and repulsive cue to RGCs, even when interacting with the same binding partner, EphB1^12,16^.

Most *in vitro* work on RGCs has been carried out on tissue culture plastics or glass. However, brain tissue is orders of magnitude softer than these materials^17^, and Eph/ephrin signalling is mechanosensitive in other contexts^18–23^. To investigate whether the discrepancies between *in vitro* and *in vivo* studies could be explained by tissue mechanics, we first cultured explants of *Xenopus* eye primordia on soft polyacrylamide substrates that mechanically mimic soft and stiff brain tissue. After one day in culture, RGCs had sent out long axons from the explants. In agreement with a previous study^24^, axon outgrowth was mechanosensitive and increased on stiffer substrates (Fig. S2a-c).

To assess the effect of substrate stiffness on Eph/ephrin signalling in RGCs, we extracted stage 35/36 eye primordia (see Fig. S1b & d-f for a schematic of developmental stages) and dissected them into thirds across their D-V axis (Fig. 1a). The developing *Xenopus* retina is characterised by a low-to-high gradient of EphB1 and EphB2 mRNA expression along its D-V axis^14^ (Fig. S1c). Therefore, the ventral thirds of the eye primordia should contain higher levels of EphB than the dorsal thirds^13,14,25,26^, and thus the response of ventral RGCs to EphB’s binding partner, ephrinB1, should be stronger than that of dorsal RGCs. This would be in agreement with previous studies in which *Xenopus*^1^, chick^8^ and mouse^9^ RGCs cultured on glass or tissue culture plastics were exposed to ephrinB proteins.

**Figure 1:**
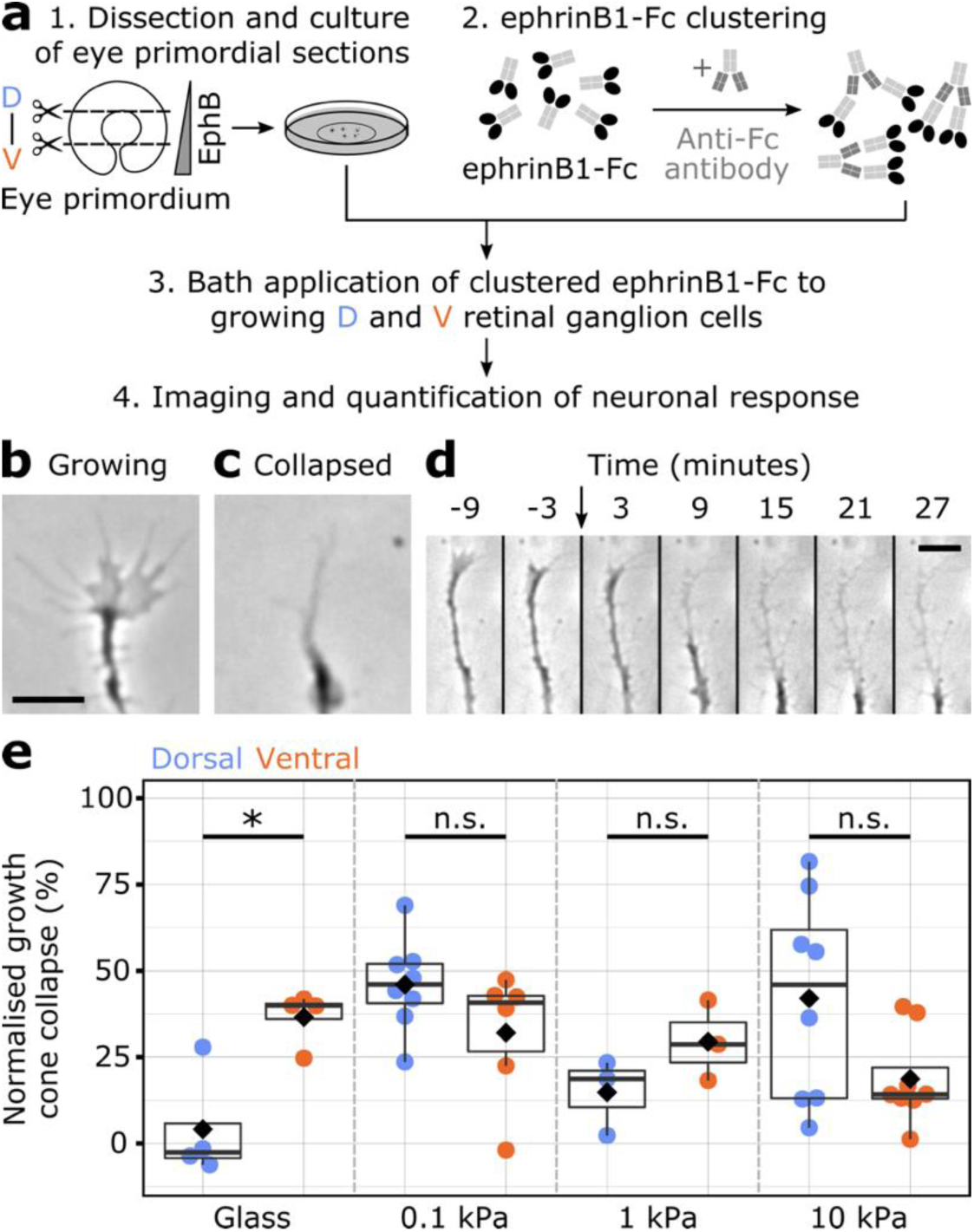
Neuronal response to ephrinB is substrate-stiffness dependent. (**a**) Schematic of the collapse assay for dorsal (D) and ventral (V) retinal ganglion cells (RGCs). Eye primordia were dissected from stage 35/36 embryos, cut into thirds, and the most dorsal and ventral sections cultured for 22-24 hours. EphB proteins are distributed in a low-to-high dorso-ventral (D-V) gradient in the retina^14^. 5 μg/mL ephrinB1-Fc was pre-clustered and applied to the growing RGCs. After 30 minutes, the cultures were fixed and the percentage of collapsed growth cones was quantified. (**b-c**) Representative images of growing and collapsed RGC growth cones. (**d**) Timelapse of a collapsing RGC growth cone after addition of ephrinB1 (black arrow). Scale bars in b-d = 10 μm. (**e**) Normalised growth cone collapse (for absolute values see Fig. S2; see Methods for details). On glass, ventral RGCs responded more to ephrinB1 than dorsal RGCs (Welch two-sample t-test; *p* = 0.0184, *t* = –3.65). On soft polyacrylamide substrates, however, the response of ventral and dorsal RGCs to ephrinB1 did not significantly differ (Welch two-sample t-tests; *p*_0.1 kPa_ = 0.156, t = 1.56; *p*_1 kPa_ = 0.189, t = –1.58; *p*_10 kPa_ = 0. 0706, t = 2.031), suggesting that Eph/ephrin signalling *in vitro* was regulated by substrate stiffness. Each point represents the mean collapse across one biological replicate (RGCs from embryos from the same *in vitro* fertilisation), the black diamond denotes the mean. For the number of growth cones *n*, see Fig. S2. *p < 0.05, n.s. = not significant.

Growth cone collapse assays, in which axon guidance activities are characterised by changes in growth cone morphology (Fig. 1b-d), were used to quantify the substrate stiffness-dependent neuronal response to 5 μg/mL ephrinB1, which was pre-clustered to allow efficient activation of EphB proteins *in vitro*^1^. On glass, as expected, growth cones of RGCs from ventral eye primordium explants collapsed significantly more than those from dorsal sections (*p* < 0.05, t = –3.65, Welch two-sample t-test; Fig. 1e & Fig. S2d). However, when explants were cultured on soft substrates with shear (storage) moduli, *G’*, of 100 Pa, 1 kPa and 10 kPa, there was no significant difference in growth cone collapse between ventral and dorsal retinal explants (all *p* > 0.05, Welch two-sample t-tests; Figs. 1e, S2e). On 0.1 kPa and 10 kPa gels, we even observed a trend for dorsal RGCs to respond more to ephrinB1 than ventral RGCs. This difference in the behaviour of neurons grown on soft substrates compared to those grown on glass was due to a significant increase in the response of dorsal RGCs to ephrinB1 on 0.1 kPa and 10 kPa gels (*p*_0.1 kPa_ < 0.05, *p*_1 kPa_ > 0.9, *p*_10 kPa_ < 0.05, one-way ANOVA followed by *post-hoc* Tukey test; Fig. S2f); the response of ventral RGCs was similar on all substrates (Fig. S2g). These results suggested that Eph/ephrin signalling in RGCs, and thus in retinotectal mapping, is mechanosensitive.

Therefore, we next characterised the stiffness of the *Xenopus* optic tectum during its innervation by RGCs, between stages 32 and 42 (Fig. S1; Fig. 2a-c), using *in vivo* AFM (Fig. 2d-f; Fig. S3a-b). To visualise tissue stiffness across the optic tectum over time, we generated averaged stiffness maps of the normalised optic tectum at three stages of innervation: pre-innervation, at innervation and post-innervation (Fig. 2g-i, see Methods for details). We found that the optic tectum was mechanically heterogeneous in both time and space. The anterior optic tectum decreased in stiffness between pre-innervation and at innervation stages (*p* < 10^-3^, Kruskal-Wallis chi-squared test followed by *post hoc* Mann Whitney U test with a Benjamini-Hochberg correction; Fig. S3c), whereas the posterior optic tectum became stiffer over time (*p*_Pre–At_ < 10^-15^, *p*_Pre–Post_ < 10^-15^, *p*_At–Post_ < 10^-3^, Kruskal-Wallis chi-squared test followed by *post hoc* Mann Whitney U test with a Benjamini-Hochberg correction; Fig. S3c).

**Figure 2:**
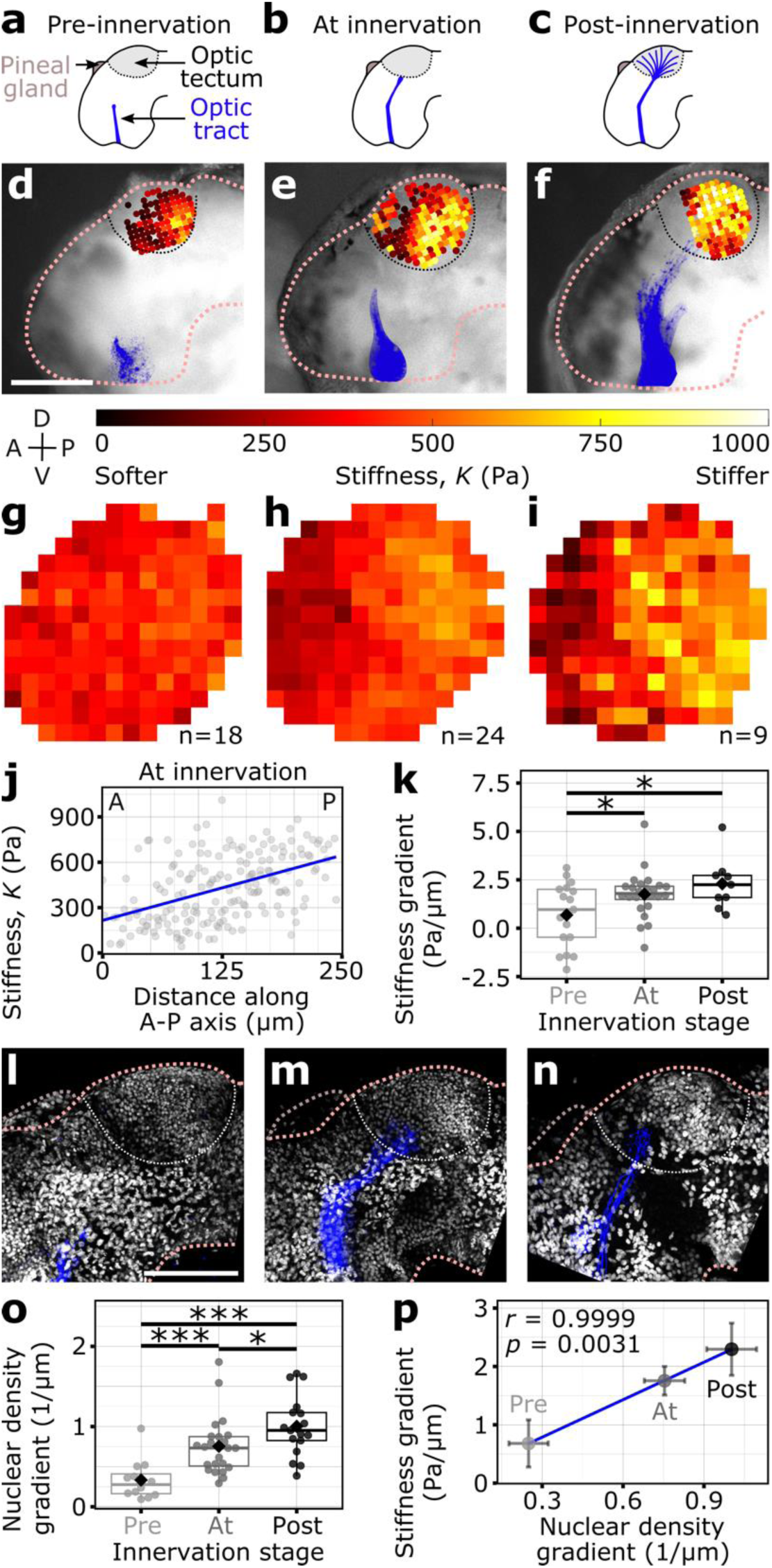
Anterior-posterior nuclear density and stiffness gradients arise in the developing optic tectum *in vivo* during innervation by RGCs. (**a-c**) Schematics of innervation stages of the *Xenopus* optic tectum (light grey) by RGCs (blue) (see Fig. S1d-f for details). (**d-f**) Example brightfield images at the stages of innervation, overlaid with AFM-based stiffness maps and DiI-labelled RGCs (blue). Anterior (A), posterior (P), dorsal (D) and ventral (V). Higher *K* values correspond to stiffer tissue. All images are scaled to the same size; scale bar = 200 μm. (**g-i**) Mean normalised scaled maps of the *Xenopus* optic tecta across stages of innervation (see Methods for details). Each pixel corresponds to the mean stiffness of that location in the tectum across *n* embryos. All heatmaps in (**d-i**) are scaled from 0 to 1000 Pa. (**j**) Quantification of a tissue stiffness gradient along the A-P axis of a stage 39 optic tectum. The slope of linear regression fit (blue) through all of the AFM measurements was extracted as the stiffness gradient value. (**k**) Pre-innervation embryos had significantly smaller stiffness gradients than those at (*p* = 0.0498) or post-innervation (*p* = 0.0199, one-way ANOVA followed by *post hoc* Tukey test), whereas there was no difference between those at innervation and post-innervation (*p* = 0.588). In all boxplots, each point represents one biological replicate *n* (*n*_Pre_ = 18, *n*_At_ = 24, *n*_Post_ = 9) and the mean is denoted by a black diamond. (**l-n**) Example maximum z-projections of nuclear density and DiI-labelled RGCs (blue) at the stages of innervation. All images are scaled to the same size; scale bar = 200 μm. (**o**) Pre-innervation embryos had smaller nuclear density gradients than those at innervation (*p* = 1.7 × 10^-5^) or post-innervation (*p* = 2.9 × 10^-6^, Kruskal-Wallis chi-squared test followed by *post hoc* Mann Whitney U test with a Benjamini-Hochberg correction). In contrast to the stiffness gradients, the nuclear density gradients were also smaller at innervation than post-innervation stages (*p* = 0.017). *n*_Pre_ = 15, *n*_At_ = 23, *n*_Post_ = 17. (**p**) A-P stiffness and nuclear density gradients correlated strongly across innervation stages (Pearson’s product-moment correlation; adjusted R^2^ = 1, Pearson’s correlation coefficient *r* = 0.999, *p* = 0.0031) and developmental stages^27^ (Fig. S3f), suggesting that changes in local nuclear density contributed to the observed A-P gradient in tissue stiffness. Each point is the mean value for the corresponding innervation stage, while the error bars denote the standard error. *n*_Pre_ = 15, *n*_At_ = 23, *n*_Post_ = 17. *p < 0.05, ***p < 0.0001. Statistically insignificant relationships are not annotated.

We then quantified the stiffness gradient across the anterior-posterior (A-P) axis by fitting a linear regression model to the measurement data from each embryo and extracting the slope of the linear fit (Fig. 2j). At younger stages (32-33/34), the stiffness gradient was negative (Fig. S3d), i.e. the optic tectum was softer posteriorly than anteriorly. In embryos at stage 35/36 and beyond, this trend was reversed and the mean stiffness gradient was greater than 0, i.e. the optic tectum was stiffer posteriorly (Fig. S3c-d). Consistent with the averaged stiffness maps of the optic tectum (Fig. 2g-i), embryos at pre-innervation stages had significantly smaller A-P stiffness gradients than those at or post-innervation (*p*_Pre–At_ < 0.05, *p*_Pre–Post_ < 0.05, one-way ANOVA followed by *post hoc* Tukey test; Fig. 2k). There was no significant difference between embryos at innervation and post-innervation stages (*p*_At–Post_ > 0.5). Thus, the stiffness gradient was not present at stage 33/34, before RGCs innervate the optic tectum, but was well established by stage 39, when RGC axons had invaded the optic tectum (Fig. S1). These stages differ by approximately 6.5 hours when developing at 22°C – 24°C^27^.

Since a highly proliferative tectal mass occupies the posterior edge of the tectal neuroepithelium^28^ and tissue stiffness in the telencephalon and diencephalon of the embryonic *Xenopus* brain is regulated by changes in cell body density on similar time scales^29^, we investigated the relationship between cell body density and tissue stiffness in the developing *Xenopus* optic tectum. Fluorescence images of cell nuclei in the *Xenopus* optic tectum at pre-innervation, at innervation and post-innervation stages indicated a change in local nuclear density over time (Fig. 2l-n). Similar to the stiffness gradient described above (Fig. 2k), the nuclear density gradient was significantly smaller at pre-innervation stages than both at and post-innervation stages (*p*_Pre–At_ < 10^-4^, *p*_Pre–Post_ < 10^-5^, Kruskal-Wallis chi-squared test followed by *post hoc* Mann Whitney U test with a Benjamini-Hochberg correction; Fig. 2o; Fig. S3e). The A-P nuclear density gradient also increased between the later stages (*p*_At–Post_ < 0.05), in contrast to the A-P stiffness gradient.

When we directly compared the stiffness and nuclear density gradients across the developing optic tectum and at different time points, we found a strong positive correlation (adjusted R^2^ = 1.00, *r* = 0.999, *p* < 0.01, Pearson’s correlation; Fig. 2p; Fig. S3f), suggesting that local cell density regulates local tissue stiffness in the *Xenopus* optic tectum.

Since Eph/ephrin signalling in RGCs innervating the optic tectum is mechanosensitive (Fig. 1), and the mechanical landscape of the optic tectum changes during the time retinotectal connections are established (Fig. 2), we next investigated the temporal development of EphB1, 2 & 4 and ephrinB1, 2 & 3 expression in the optic tectum using hybridisation chain reaction (HCR) RNA-FISH (Fig. 3; Fig. S4).

**Figure 3:**
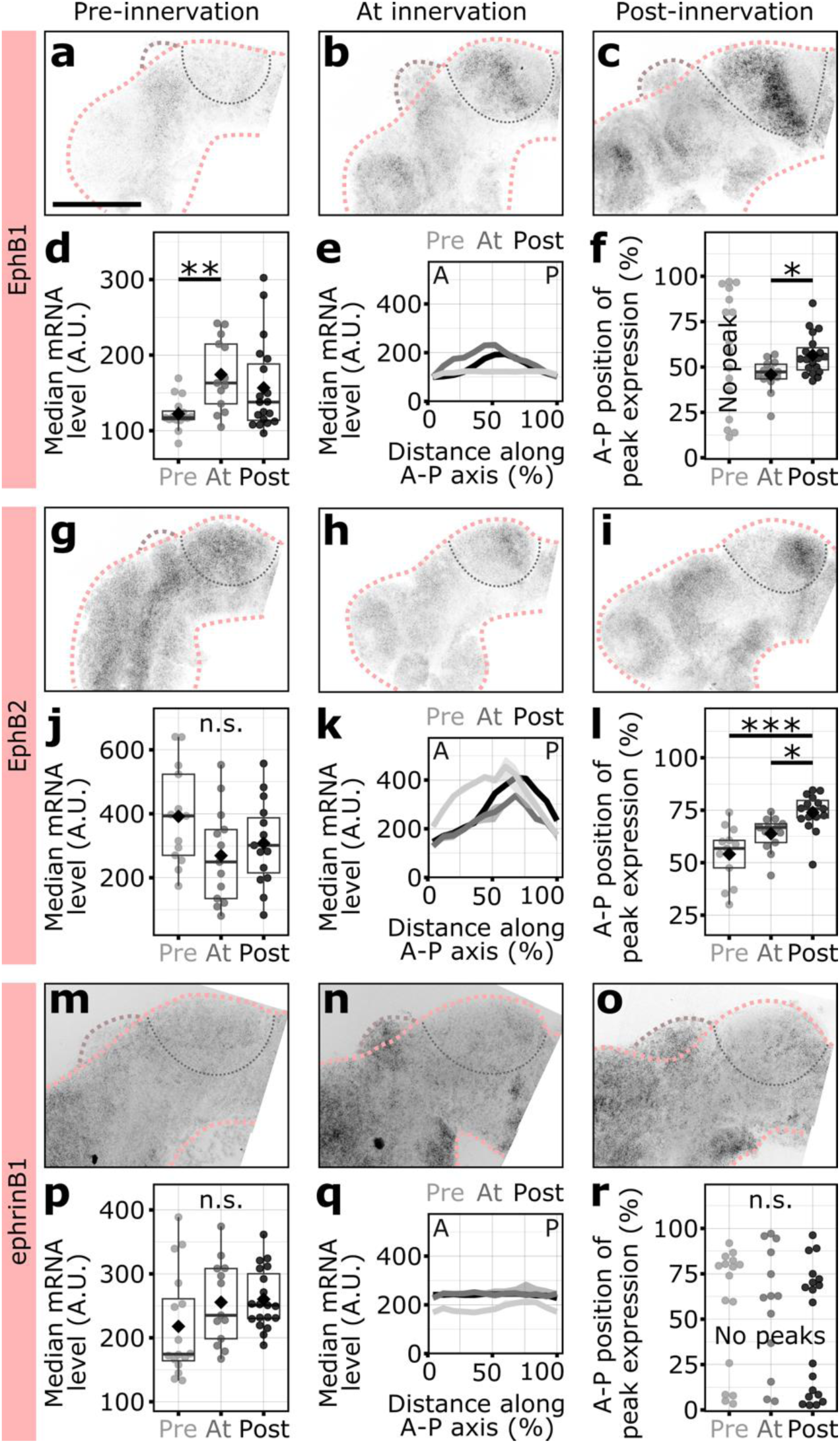
EphB/ephrinB expression along the anterior-posterior axis of the optic tectum during innervation by RGCs. Representative images of EphB1 (**a-c**), EphB2 (**g-i**) and ephrinB1 (**m-o**) mRNA expression in the optic tectum during innervation by RGCs (see Fig. S1 for details). Scale bar = 200 μm. Grey values scaled within each gene. (**d, j, p**) Quantification of median mRNA levels for EphB1 (**d**), EphB2 (**j**) and ephrinB1 (**p**). EphB1 expression increased over time (*p*_Pre–At_ = 0.0017, *p*_Pre–Post_ = 0.182, *p*_At–Post_ = 0.182, Kruskal-Wallis chi-squared test followed by *post hoc* Mann Whitney U test with a Benjamini-Hochberg correction), whereas EphB2 (*p* = 0.1, F-value = 2.444, one-way ANOVA) and ephrinB1 (*p* = 0.076, χ^2^ = 5.1443, Kruskal-Wallis chi-squared test) expression did not change. (**e, f, k, l, q, r**) Quantification of anterior-posterior peaks in mRNA levels of EphB1 (**e, f**), EphB2 (**k, l**) and ephrinB1 (**q, r**) at the different stages. The peak of EphB1 expression shifted posteriorly past the mid-tectum during development (*p* = 0.00544, t = –3.0024, Welch two-sample t-test), whereas that of EphB2 shifted from the mid-to the posterior tectum during innervation (*p*_Pre–At_ = 0.0542, *p*_Pre–Post_ = 0.0000302, *p*_At–Post_ = 0.0350, one-way ANOVA followed by *post-hoc* Tukey test). EphrinB1 expression was homogeneous along the A-P axis at all stages (*p* = 0.428, F-value = 0.865, one-way ANOVA). Negative controls (HCR RNA-FISH hairpins only) are shown in Fig. S4. In all boxplots, each point represents one biological replicate (brain) from three different *in vitro* fertilisations. The mean is denoted by a black diamond. In each line profile, the median grey value of every 20 µm is indicated by a line and a ribbon denotes the 95% confidence interval. *p < 0.05, **p < 0.01, ***p < 0.001, n.s. = not significant. Statistically insignificant relationships are not annotated in **d**, **f**, and **l**.

EphB1 mRNA levels increased significantly in the optic tectum upon innervation by RGC axons (*p*_Pre–At_ = 0.0017, Kruskal-Wallis chi-squared test followed by *post hoc* Mann Whitney U test with a Benjamini-Hochberg correction; Fig. 3a-d), and peak expression shifted posteriorly past the mid-tectum between at innervation and post-innervation stages (*p* < 0.01, Welch two-sample t-test; Fig. 3e, f). Although EphB2 mRNA levels did not change significantly throughout the entire tectum (*p* = 0.1, one-way ANOVA; Fig. 3g-j), the peak expression of EphB2 shifted from the mid-to the posterior tectum during innervation (*p*_Pre–Post_ < 10^-4^, *p*_At–Post_ < 0.05, one-way ANOVA followed by *post-hoc* Tukey test; Fig. 3k, l).

EphrinB3 showed a similar shift in expression peaks from the mid-to the posterior tectum between pre-innervation and at innervation stages (*p*_Pre–At_ = 0.0005, Kruskal-Wallis chi-squared test followed by post hoc Mann Whitney U test with a Benjamini-Hochberg correction; Fig. S4l-o). In comparison, EphB4, ephrinB1 and ephrinB2 mRNA levels remained consistently low with no distinct patterns throughout innervation (Fig. 3m-r, *p* > 0.05, Kruskal-Wallis chi-squared test; Fig. S4h-k). Negative controls were used to confirm low levels of non-specific binding of fluorescent hairpins to *Xenopus* brain tissue at these stages (Fig. S4p-w). Together, these experiments showed that EphB and ephrinB family members have different expression patterns across the A-P axis of the optic tectum during innervation by RGC axons.

To further investigate the relationship between tissue stiffness and Eph/ephrin expression patterns, we next conducted a correlation analysis. We first generated averaged expression maps of the normalised optic tectum over time for each gene, and then plotted the mean mRNA level and stiffness values for each coordinate of the normalised tectum against each other (Fig. S5, see Methods for details). With Pearson’s correlation coefficients *r* ∼ 0.8, the expression patterns of EphB2, EphB4 and ephrinB3 strongly correlated with the stiffness patterns at post-innervation stages, whereas EphB1 and ephrinB1 expression weakly correlated with tissue stiffness (*r* ∼ 0.3), and only ephrinB2 expression did not correlate with tissue stiffness (*r* ∼ 0) (Fig. S5, see Table S1 for details). This suggested that tissue stiffness may regulate the expression of Ephs and ephrins in the developing brain.

To test this hypothesis, we performed two assays to alter the mechanics of the *Xenopus* brain. First, we embedded dissected brains at stage 33/34, before the stiffness gradient across the optic tectum arises (Fig. S3d), in 3D hydrogels of G’ 40 Pa (soft) and 450 Pa (stiff) for 24 hours (Fig. 4a). Because the response of RGCs to ephrinB1 was mechanosensitive (Figs. 1, S2), we quantified the mRNA expression of ephrinB1, and EphB1 and EphB2, both of which are high-affinity binding partners of ephrinB1^30,31^ in the optic tectum. If Eph/ephrin gene expression is indeed regulated by the mechanical properties of the environment, mRNA levels should be higher in tissue embedded in the stiffer gels. Accordingly, the median mRNA expression levels of EphB1 and EphB2 were significantly higher in the stiff than in the soft condition (*p*_EphB1_ < 0.01, t_EphB1_ = –3.17; *p*_EphB2_ = 0.05, t_EphB2_ = –2.50, Welch two-sample t-tests; Fig. 4b-g) but independent of the stiffness of the environment in case of ephrinB1 (*p* > 0.05, t = –1.94, Welch two-sample t-test; Fig. 4h-j). These data were consistent with our previous findings that the level of EphB2 expression, but not that of ephrinB1, was strongly correlated with tissue stiffness (Fig. S5b, d).

**Figure 4:**
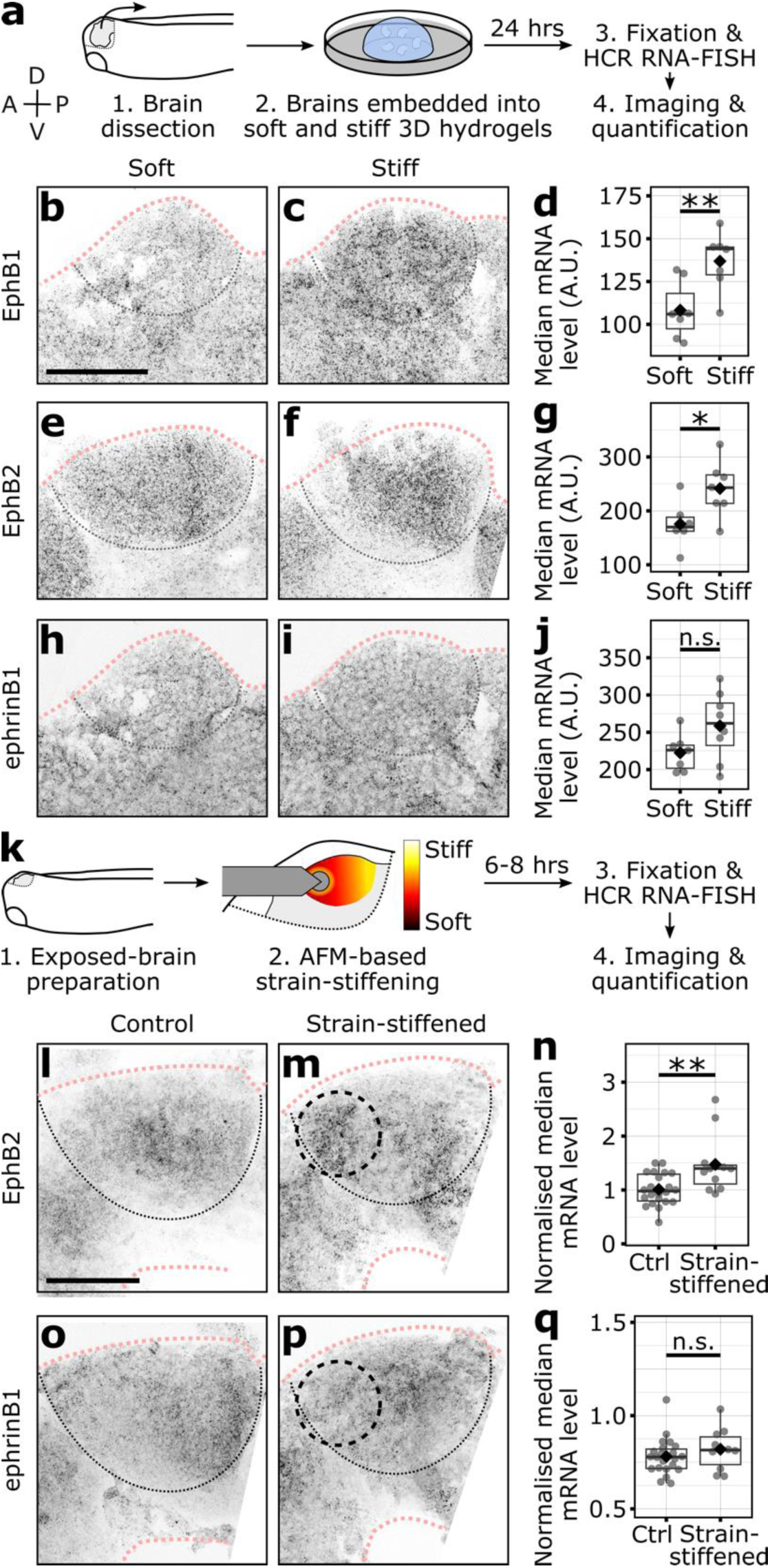
Mechanical perturbations of the optic tectum lead to changes in the expression of EphB1 and EphB2, but not ephrinB1. (**a**) Schematic of mechanical perturbation of the brain environment *ex vivo*. Stage 33/34 *Xenopus* brains were embedded in soft (G’ = 40 Pa) and stiff (G’ = 450 Pa) 3D hydrogels for 24 hours and mRNA expression was quantified. Anterior (A), posterior (P), dorsal (D) and ventral (V). (**b, c, e, f, h, i**) Representative images of EphB1 (**b-c**), EphB2 (**e-f**) and ephrinB1 (**h-i**) mRNA expression after exposure to soft and stiff 3D hydrogels. Image intensity scaled within each gene. (**d, g, j**) HCR quantification. EphB1 (**d**) and EphB2 (**g**) mRNA levels were significantly higher in optic tecta in stiff if compared to soft hydrogels (*p*_EphB1_ = 0.00803, t_EphB1_ = –3.173; *p*_EphB2_ = 0.0294, t_EphB2_ = –2.502, Welch two sample t-tests), whereas there was no difference in ephrinB1 mRNA levels (**j**) (*p* = 0.0786, t = –1.938). (**k**) Schematic of local AFM-based strain-stiffening of optic tecta *in vivo*. A constant force of 30 nN was applied to the anterior, softer region of the optic tectum of stage 33/34 brains for 6-8 hours using a ∼90 μm spheroidal probe, leading to local tissue stiffening^24,35^. The embryo was then fixed and mRNA expression quantified. (**l, m, o, p**) Representative images of EphB2 (**l-m**) and ephrinB1 (**o-p**) mRNA expression for control (**l, o**) and strain-stiffened (**m, p**) embryos. The dashed black circles indicate the position of the AFM bead during strain-stiffening. Image intensity is scaled so the posterior half of the optic tectum is of similar intensity for each gene in both conditions. (**n, q**) HCR quantification. (**n**) EphB2 mRNA levels were significantly higher in the strain-stiffened regions than in the corresponding region in control brains (*p* = 0.007, W = 49.5, Mann-Whitney U test), while there was no difference in ephrinB1 expression (*p* = 0.255, W = 81.5). These data indicate that mechanical perturbations of brain tissue were sufficient to alter EphB2, but not ephrinB1, expression in the *Xenopus* optic tectum. In all boxplots, each point represents one biological replicate, obtained from three and four different *in vitro* fertilisations for the 3D hydrogel and strain-stiffening experiments, respectively. The black diamond denotes the mean. Scale bars = 100 μm.

To corroborate these results, we directly altered local brain mechanics *in vivo*. Brain tissue stiffens under compression^24,32–34^, enabling local and precise stiffening of brain tissue by AFM^24,35^. To stiffen the softer, anterior region of the optic tectum in stage 33/34 embryos, we locally applied a constant compressive force of 30 nN for 6-8 hours at 18°C using a ∼90 µm spheroidal AFM probe. Embryos were then fixed and mRNA expression quantified using HCR RNA-FISH (Fig. 4k). Since the expression levels of EphB2, but not ephrinB1, were altered by global mechanical perturbation *ex vivo* (Fig. 4e-j), and EphB2 expression correlated strongly with tissue stiffness *in vivo* (Fig. S5b), we focused on these two candidates. While strain-stiffening of brain tissue had no effect on ephrinB1 mRNA levels (*p* > 0.2, W = 81.5, Mann-Whitney U test; Fig. 4o-q), EphB2 mRNA levels were significantly higher in the strain-stiffened condition compared to the control (*p* < 0.01, W = 49.5, Mann-Whitney U test; Fig. 4l-n). Together, these experiments showed that, *in vivo*, an increase in brain tissue stiffness, which occurs in the posterior part of the optic tectum during development (Figs. 2i, S3c), is sufficient to increase EphB2, but not ephrinB1, expression levels.

## Discussion

We have shown that both the response of RGCs to Ephs and ephrins and the expression of Ephs and ephrins in the developing optic tectum are mechanosensitive. We found that the *Xenopus* optic tectum becomes mechanically and chemically heterogeneous as it is innervated by RGC axons, and that these mechanical and chemical changes may be linked. We found a strong correlation between tissue stiffness and the expression of EphB2, EphB4 and ephrinB3, and changes in tissue mechanics led to changes in EphB2 expression *ex vivo* as well as *in vivo*. Hence, the A-P tectal stiffness gradient (Figs. 2, S3), which precedes the changes in EphB2 gene expression by a few hours (Fig. 3l), likely contributes to the observed EphB2 patterns along the A-P axis of the optic tectum during development.

Based on coronal cross-sections and *in vivo* mutagenesis, EphB/ephrinB signalling in the *Xenopus* retinotectal system has so far been implicated mainly in the mapping of RGC axons along the D-V axis of the optic tectum^13,14^. Here, we identified differential expression patterns of EphB1, EphB2, EphB4 and ephrinB3 along the A-P axis that emerge between stages 37/38 and 40, during the innervation of the optic tectum by RGC axons (Fig. 3; Fig. S4). The role of EphB/ephrinB signalling in the establishment of the D-V axis of the retinotectal projection was supported by *in vitro* experiments on glass substrates, where RGCs from the ventral retina responded more strongly to ephrinB than dorsal RGCs^1,8,9,14^. We replicated these established findings on glass, but subsequently found that dorsal and ventral RGCs respond similarly to ephrinB1 on substrates with a stiffness similar to that found in the developing *Xenopus* optic tectum (Fig. 1e). In line with our interpretation that substrate stiffness regulates neuronal responses to ephrinB, two early *in vitro* studies performed on monolayers of soft tectal cells and carpets of tectal membrane fragments of ventral or dorsal origin also found that dorsal and ventral chick RGCs responded similarly to both^7,11^, despite the fact that the chick dorsal tectum contains high levels of ephrinB^36^. The dependence of Eph/ephrin signalling on the mechanical environment described here may thus contribute to the discrepancies between previous *in vivo* studies^12,13^ and *in vitro* experiments conducted on glass or tissue culture plastics^1,8,9^, which are much stiffer than neural tissue.

Prior to the arrival of RGC axons, an A-P stiffness gradient emerged across the optic tectum, which peaked at ∼2 Pa / µm at stage 35/36 and remained stable thereafter (Fig. 2; Fig. S3d). This stiffness gradient positively correlated both temporally and spatially with a nuclear density gradient across the optic tectum (Fig. 2p), suggesting that local cell density may drive changes in local tissue stiffness in the developing *Xenopus* optic tectum, as previously shown for the *Xenopus* embryonic telencephalon and diencephalon^29^.

Using mechanical perturbations, we found that the stiffness gradient is not just an epiphenomenon of the pattern of tectal cell proliferation. Instead, it regulates the expression of EphBs and ephrinBs (Fig. 4). Consistent with these experiments and our *in vivo* tissue stiffness and mRNA expression correlation analysis (Fig. S5b,c,f, Table S1) showing that stiffer tissues drive the expression of EphB2, EphB4 and ephrinB3, during *Xenopus* gastrulation, these same three genes are preferentially expressed in the ectoderm rather than in the mesoderm^37^, with the ectoderm being stiffer than the mesoderm^38^.

Our mechanical perturbation experiments (Fig. 4), together with the mRNA expression data (Fig. 3; Fig. S4), also suggested that there may be differences in the mechanosensitive regulation of the expression of different Ephs and ephrins. For example, the correlation between tissue stiffness and gene expression was much stronger for EphB2 than for ephrinB1 (Fig. S5b,d), and in contrast to EphB2, ephrinB1 was not significantly upregulated after strain-stiffening of the optic tectum (Fig. 4h-j, o-q). The mechanotransduction pathways associated with Eph/ephrin signalling are not yet understood, but the relationship between chemical and mechanical cues is clearly reciprocal.

The upstream regulation of Eph/ephrin signalling is currently not fully understood. However, Ephs and ephrins are transmembrane proteins and their activity may not only be regulated by chemical signals but also directly be influenced by the physical properties of the plasma membrane. For instance, the tension of the plasma membrane and its linkage to the cytoskeleton impact processes such as receptor clustering and endocytosis^39–41^ that are essential for Eph/ephrin signalling^2^. Substrate stiffness has been shown to affect endocytosis^42–44^, and membrane tension is very different on soft substrates compared to glass^45^. On the other hand, Eph/ephrin signalling also affects the levels of membrane-actin cortex linkers^46^, cortical tension^47^ and the arrangement of adhesion molecules^48,49^. This in turn affects how RGCs sense their environment and interact with other cells, leading to a feedback mechanism^50^.

To date, most studies of signalling pathways in biology investigate either chemical or mechanical signals. At present, there is little evidence for a close link between the two signalling modalities^51^. Here we show that the presence and abundance of the receptor protein tyrosine kinase, Eph, and its binding partner, ephrin, are regulated by tissue mechanics. Untangling the interplay between mechanical and chemical cues will not be trivial, but is necessary to understand and predict how the brain and other organ systems develop *in vivo*. As Eph/ephrin signalling is a highly conserved signalling family, understanding how it is modulated by tissue mechanics has vast implications for the general understanding not only of retinotectal mapping, but also of many other crucial processes across a wide range of organ systems and species, including bone homeostasis^3^, spinal cord injury^17,52^ and epithelial-to-mesenchymal transitions in development and cancer^22^.

## Methods

Detailed methods are available in the online version of the paper, including the associated code and references.

## Data availability

The datasets generated during and/or analysed during the current study are available from the corresponding authors on reasonable request.

## Supporting information

Sipkova Supplement

## Acknowledgements

We thank A. Winkel for AFM measurements of hydrogels, J.M. Becker and A. Winkel for providing and optimising AFM analysis code, S. Mukherjee for establishing the HCR RNA-FISH protocol in Xenopus brain tissue, N. Gampl for establishing 3D collagen-based hydrogel preparation and culture, and the rest of the Franze and Holt labs for discussions and extensive help in the lab. This work was supported by a Wellcome Trust scholarship 215156/Z/18/Z (J.S.), a European Research Council Consolidator Award 772426 (K.F.), the German Research Foundation (DFG) projects 460333672 CRC1540 EBM and 270949263 GRK2162 (K.F.) and an Alexander von Humboldt Professorship (Alexander von Humboldt Foundation) (K.F.).

## Author contributions

J.S. and K.F. conceived the project and designed the research; J.S. performed the experiments and analysed the data; both authors discussed the data; J.S. wrote the original draft of the manuscript; J.S. and K.F. revised the manuscript to the final version.

