## Supplementary material for "Eph/ephrin signalling in the developing brain is regulated by tissue stiffness": Sipkova Supplement

### Online Methods

All reagents were obtained from Sigma-Aldrich unless otherwise specified.

#### Xenopus laevis husbandry

All animal experiments were approved by the Ethical Review Committee of the University of Cambridge and complied with Home Office guidelines. *Xenopus laevis* embryos were fertilised *in vitro*, raised in 0.1× Modified Barth's Saline (MBS) or Marc's Modified Ringer's (MMR) solution at 14–16°C, and staged according to Nieuwkoop and Faber[1].

Embryos were classified into three categories based on both the general morphology[1] and the extent of optic tract development[2]: the optic tract had not yet reached or had just reached the characteristic bend in the diencephalon (stages 32-35/36, pre-innervation), the optic tract was just crossing the boundary of the optic tectum (stages 37/38-40, at innervation), and the RGC axons were beginning to spread out in the optic tectum (stages 41-44, post-innervation) (Fig. S1d-f).

Unless otherwise stated, prior to fixation and dissection all embryos were anaesthetised in 1:1 MS222 solution (0.04% w/v tricaine methanesulfonate salt (MS222) and 1× Penicillin-Streptomycin-Amphotericin B, PSF (Lonza), in 1× MBS/MMR, pH 7.6) and 0.1× MBS/MMR (fixation) or culture medium (dissection; 60% Leibovitz L15 medium, 1× PSF, pH 7.6-7.8). All dissections were carried out by pinning down embryos in a Sylgard 184-coated (Dow Corning) dissection dish with bent 0.2 mm minutien insect pins (Austerlitz), and dissecting embryos using 0.15 and 0.1 mm minutien insect pins (Austerlitz) in pin holders.

**Whole-mount *Xenopus* brain Hybridisation Chain Reaction RNA fluorescence *in situ* hybridisation (RNA-FISH)**

A modified Hybridisation Chain Reaction (HCR) RNA-FISH protocol (Molecular Instruments)[3] based on Aztekin et al., 2021[4] and developed by Pillai, Mukherjee et al, 2024[5], was performed on whole-mount *Xenopus* brains at stages of interest. We are extremely grateful to Jakub Sedzinski (The Danish Stem Cell Centre, University of Copenhagen, Denmark) for guidance with troubleshooting and optimising the protocol.

Embryos were fixed in modified Carnoy's fixative[6] (6 vol 100% ethanol, 3 vol 37% formaldehyde, 1 vol glacial acetic acid) for 4 hours at room temperature, or overnight at 4°C. After rinsing with 70% ethanol in PBST (PBS and 0.1% Tween-20) for 5-10 min, embryos were dehydrated into 100% ethanol in three 30 min washes, and stored at -20°C.

Embryos were rehydrated in 75%, 50% and 25% ethanol in PBST in sequential 15 min washes, and then washed three times in PBST for 15 min each. Brains were dissected out of the embryos and transferred into a 4-well plate (Fisher Scientific) with PBST. Dissected brains were then bleached in bleaching solution (5% formamide, 2.5% 20× SSC and 28% $H_2O_2$ diluted in distilled water) for 45 min under a strong light source. After bleaching, the brains were gently rinsed in water, and then washed in PBST three times 10 min each. They were subsequently treated with 5 µg/ml Proteinase K diluted in PBST for 10 min at room temperature, then gently washed three times 5 min each in PBST. Brains were fixed again for 20 min with 3.7% formaldehyde in PBST and washed 5 times for 5 min in PBST.

The hybridisation and amplification steps followed the protocol from Aztekin et al., 2021[4]. Briefly, samples were washed in 1 mL pre-heated probe wash buffer for 5 min at room temperature, and then in 500 µl pre-heated hybridisation buffer for 30 min at 37°C. In parallel, the probe solution was prepared by diluting Molecular Instrument custom probes in 500 µL hybridisation buffer, to a final concentration of 12 nM. The probe solution was pre-heated at 37°C for 30 min, and subsequently the hybridisation buffer was replaced with the probe solution and samples left overnight at 37°C.

Samples were washed thoroughly after hybridisation: twice for 30 min in pre-heated wash buffer, once for 5 min in 50% 5× SSCT/50% wash buffer, and twice for 20 min in 100% 5× SSCT, all at room temperature. In parallel, the appropriate hairpins were heated to 95°C for 90 sec and left for 30 min in the dark at room temperature. The amplification solution was prepared by diluting the hairpins in 500 µL amplification buffer, to a final concentration of 24 nM. The samples were pre-amplified with 1 mL amplification buffer at room temperature for 10 min, and then incubated with the amplification solution with hairpins overnight at room temperature, in the dark. Samples were subsequently washed twice for 30 min in 5× SSCT and stored in 1× PBS at 4°C until mounting and imaging.

50 µm z-stacks were acquired on a confocal microscope in the Cambridge Advanced Imaging Centre (SP8, Leica Microsystems; 40×/1.3 oil; z-step size = 2 µm). Multiple tiles were acquired for each whole-mount brain, ensuring that the optic tectum fit within one tile so stitching artefacts did not interfere with subsequent analysis. Tile stitching was performed within the Leica LAS X software immediately after image acquisition. Image analysis was

performed on the outer-most 16 µm of the brain, through which the RGC axons initially grow[7]. Representative images shown are average z-projections of the entire z-stack.

#### Visualising the optic tract by DiI labelling

DiI was used for the lipophilic labelling of the entire optic tract as previously described[8,9]. Embryos at the developmental stage of interest, or after AFM, were fixed using 4% PFA in 1× PBS for 1.5-2 hours at room temperature or overnight at 4°C. After a brief wash in 1× PBS, embryos were pinned down in a Sylgard 184-coated (Dow Corning) in 1× PBS. For embryos which had been measured through AFM, the embryo was pinned with the side containing the intact eye facing up, to allow for labelling of the RGCs on the exposed side of the brain. A working solution of DiI (1,1'-Dioctadecyl-3,3,3',3'-Tetramethylindocarbocyanine Perchlorate; Molecular Probes) dissolved in 100% ethanol was heated for a 2-3 min at 65°C. The solution was loaded into injection pipettes pulled from 1.0mm OD × 0.78 mm ID capillaries (without filament; Harvard Apparatus). Using a Femtojet 4X microinjector (Eppendorf), DiI was injected into the eye, at the boundary between the lens and the retinal pigmented epithelium (Fig. S3b). Embryos were left protected from light in 1× PBS for 24-28 hours at room temperature, allowing the DiI to diffuse along the length of the RGC axons. The brains were then dissected out and either directly mounted in 1× PBS in the lateral view for imaging or stained for nuclear density with DAPI. Those directly imaged were imaged using a Zeiss AxioObserver.A1 inverted microscope with an EC Plan-Neofluar 10×/NA 0.3 Ph1 M27 air objective (Zeiss), using an sCMOS camera (Andor Zyla 4.2). Both phase contrast and epifluorescence images were acquired. For the latter, a metal halide lamp (Zeiss) and modified TRITC filter set (Zeiss, No. 45; absorption 560/40 (BP)/emission 630/75 (BP)) were used to visualise the optic tract.

#### Labelling nuclei in whole-mount brains

After HCR RNA-FISH or DiI injection, whole-mount *Xenopus* brains were incubated in 1 µg/mL DAPI (4',6-diamidino-2-phenylindole, Santa Cruz Biotechnology) diluted in PBS for 10-15 min. Brains were washed in 1× PBS for 10 min and subsequently mounted in 1× PBS in the lateral view for imaging. Z-stacks were acquired on a confocal microscope in the Cambridge Advanced Imaging Centre (SP8, Leica Microsystems; 20×/0.75 air; z-step size = 1 µm). For the whole-mount brains stained after DiI injection, z-stacks spanned the entire brain (usually 40-45 µm), and were subsequently used for nuclear density analysis.

#### Atomic force microscopy (AFM)

The health and viability of embryos throughout AFM measurements and local strain-stiffening was assessed by the presence of a visible heartbeat[10] and integrity of the skin. A brightfield overview image of the entire embryo was taken prior to and after each AFM experiment and used for comparison.

##### *Exposed brain preparation*

A modified exposed brain protocol was used to allow for direct AFM measurements of the brain surface, as previously described[9,11,12]. Embryos were anaesthetised in 0.04% w/v MS222, 1× PSF, in 1.3× MBS, pH 7.6, henceforth referred to as exposed brain medium, and the eye, epidermis and dura on one side was removed, exposing the optic tectum. The higher salt concentration slows skin regrowth, allowing for AFM measurements to be taken over an extended period. Embryos were subsequently transferred to and immobilised in Sylgard 184-coated (Dow Corning) 35-mm Petri dishes (World Precision Instruments).

*Cantilever and instrument preparation*

Cantilevers were calibrated and mounted as described previously[9,11]. Briefly, tipless silicon cantilevers (Arrow-TL1; NanoWorld) were calibrated using the contact-based thermal noise method[13] in a built-in JPK software (JPK Instruments, Bruker). Cantilevers with spring constants *k* of 0.01-0.04 N/m were selected for measurements of the *Xenopus* brain and compliant hydrogel substrates. Monodisperse spherical polystyrene beads (microParticles GmbH) with a diameter of 37.28 µm were attached to the cantilever ends using M-Bond 610 heat-curing glue (MicroMeasurements)[14,15]. The probe size was chosen according to the size of the sample and scale of the measurement. Average indentations of approximately 3-7 µm probed the environment of the RGC axon growth cones, which grow within the top 10 µm of the tissue[7]. As a result, contact radii were approximately 7.3-10.7 µm[16–18].

Prior to sample measurement, a cantilever was mounted on a CellHesion-200 AFM head (JPK Instruments, Bruker), set up on motorised stage of a Zeiss AxioObserver.A1 inverted microscope. To allow for upright imaging of the *Xenopus* brain during AFM measurements, a modified Zeiss AxioZoom V.16 system was mounted on a custom-built support arm above the AFM head as described in Thompson, Pillai et al., 2019[9].

In vivo *AFM*

AFM on live embryos was performed as previously described[9,11]. Briefly, beaded cantilevers were first calibrated in exposed brain medium to determine the deflection sensitivity. After exposed brain preparation and mounting the sample, an overview image of the brain was taken using a PlanApo Z 0.5×/0.125 objective (working distance 114 mm,

Zeiss) and sCMOS camera (Andor Zyla 4.2). Three reference points and their corresponding motorised stage coordinates were selected in the overview image. Using a custom-written MATLAB script (courtesy of Julia Becker and Alex Winkel, University of Cambridge), the region of interest was selected, and a grid of measurement coordinates was generated. Indentations were spaced by 15 µm, which is of appropriate scale to how much growth cones can sense *in vivo*[7]. Custom-written Python scripts (courtesy of Amelia J. Thompson, Julia Becker and Alex Winkel, University of Cambridge) read the list of coordinates into the JPK software and measurements were performed, generating a 2D 'stiffness map'. For each measurement, an image of the cantilever and sample was recorded, and a force curve taken (approach speed: 20 µm/s, force: 10 nN, sample rate: 1000 Hz). The cantilever was then retracted by 100 µm and the stage moved to the next position. Each map took approximately 30-50 min to record. Since tissue stiffness in the *Xenopus* optic tectum can rapidly change[9], measurements were carried out in different directions across biological replicates within one experiment to ensure that the temporal aspect of the measurement did not bias one side to always being older. After AFM mapping, embryos were fixed in 4% PFA, and the optic tract labelled with DiI to help determine the stage of RGC innervation of the optic tectum (Fig. S3a).

*Strain-stiffening using AFM*

Local strain stiffening experiments were performed as previously described[11]. Embryos were treated the same way as for *in vivo* AFM, apart from the settings of AFM application. Briefly, stage 33/34 embryos were anaesthetised, and one side of the optic tectum exposed in exposed brain medium. Overview brightfield images of the entire embryo and the brain were taken using the same system as described above. Tipless silicon cantilevers (Arrow-

TL1; NanoWorld) with attached polystyrene beads (microParticles GmbH) with a diameter of 89.3 µm and spring constants *k* of 0.2-0.35 N/m were used to apply a constant force of 30 nN to the anterior region of the optic tectum for 6-8 hours at 18°C. Control embryos were similarly pinned down in dishes with their brains exposed but did not undergo AFM application. After strain stiffening, both strain-stiffened and control embryos were immediately fixed in modified Carnoy's fixative[6] and stored at -20°C for HCR RNA-FISH (Fig. 4k). After HCR RNA-FISH, the overview brightfield image taken during strain-stiffening was overlaid on the fluorescence images of EphB/ephrinB expression using the pineal gland and posterior boundary of the optic tectum as landmarks. The position of the AFM bead and therefore the region which was locally stiffened was thus determined in the strain-stiffened condition. Importantly, only the brightfield and DAPI images taken along with the fluorescence images, and not the images containing EphB/ephrinB mRNA signal, were used to determine the location of the bead, eliminating possible bias. For quantification, the median mRNA level in the strain-stiffened region of the optic tectum or the corresponding region in the control condition was normalised by the median mRNA level in a region of the diencephalon with low expression to account for variation between different biological replicates.

**AFM data analysis**

*Processing of raw AFM data*

Custom-written MATLAB scripts were used to calculate the reduced apparent elastic modulus, *K*, from force-distance curves generated from stiffness measurements (modified from Christ et al., 2010[19], Koser et al., 2016[11], and Thompson, Pillai et al., 2019[9] by Julia

Becker and Alex Winkel, University of Cambridge; https://github.com/FranzeLab/AFM-data-analysis-and-processing/tree/master/JuliaBeckerThesis).

Briefly, the raw AFM data were fit to the Hertz model for a spherical indenter,

$$F = \frac{4}{3}\frac{E}{1-\nu^2}\delta^{\frac{3}{2}}\sqrt{R}$$

where **F i**s the known applied force in N, ***E*** the Young's modulus in Pa, **ν** the Poisson ratio, δ the measured indentation depth, and **R** the radius of the indenter[20,21]. **ν** is an intrinsic property of the material describing its change in length upon changes in its cross-section and must be measured or assumed. Since the few reported experimental measurements of cells and tissues vary, **ν** is often assumed to be 0.5, due to the water content of cells and tissues rendering them incompressible[22].

However, drawing on Hooke's law, the Hertz model can also be expressed as

$$F = \frac{4}{3}K\delta^{\frac{3}{2}}\sqrt{R}$$

where ***K*** is the reduced apparent elastic modulus and is equivalent to $E/(1-\nu\hat{}2\ )$.

In this equation, all parameters apart from *K* are known or measured using the AFM. Thus, expressing tissue stiffness as *K*, as opposed to E, allows for ν to stay undefined.

Force-distance curves were analysed at the maximum applied force, which was typically 10 nN. Measurement points where the raw AFM data could not be analysed were excluded. This included force-distance curves which either could not be fit with a linear fit at the baseline region, due to for example noise, or could not be fit with the Hertz model in the indentation region.

The stiffness data from each experiment were arranged into a 2D array (x, y) to be used for both subsequent analysis in R and visualisation in MATLAB using a custom-written script by Julia Becker, University of Cambridge (https://github.com/FranzeLab/AFM-data-analysis-and-processing/tree/master/JuliaBeckerThesis/c_After%20AFM%20measurements). For visualisation of AFM stiffness maps, values of *K* were converted into an 8-bit scale and visualised as heatmaps using the 'hot' pre-set MATLAB colourmap.

*Quantifying stiffness across the optic tectum*

The quantification of the stiffness across the optic tectum was conducted using custom-written MATLAB and R scripts (https://github.com/FranzeLab/AFM-data-analysis-and-processing/blob/master/SipkovaPaper2024/AFM_grid_tectum_analysis_FIRST.m) .

To carry out quantification across different biological replicates, the AFM maps were normalised in *x* and *y*. A brightfield image associated with each sample was first used to rotate each embryo so that the A-P axis was horizontal, and anterior was facing left. Then, the brightfield image was used to create an outline of the optic tectum, and the AFM measurement values found within the outlined region were extracted. The raw measurement coordinates of the corresponding measurement values were subsequently rescaled to adjust for the previously established angle of rotation, and then normalised from 0 to 1 to allow comparison between biological replicates with varying tectum sizes. In R, the normalised *x* and *y* coordinates, along with the associated tissue stiffness values, were then analysed using linear regression.

*Generating mean normalised stiffness maps of the optic tectum*

Using the output from the MATLAB script described above, another MATLAB script was used to generate mean normalised maps of the optic tectum at different stages of innervation (https://github.com/FranzeLab/AFM-data-analysis-and-processing/blob/master/SipkovaPaper2024/Avg_tectum_map_SECOND.m). This allows for visual comparison of general trends in the optic tectum across time and can be easily applied to other AFM datasets. The data from all normalised biological replicates was first plotted on a dummy optic tectum. Then, a boundary was generated around the datapoints, and a grid was created within this boundary. The coordinates of each stiffness value were reassigned to the closest grid point, and subsequently the mean stiffness value at each grid point was calculated. This dataset was then plotted in a heatmap, to visualise the mean normalised optic tectum across the input data.

**Preparation of hydrogels for cell culture**

*Fabrication of 2D polyacrylamide gels*

Polyacrylamide (PAA) gel substrates of 100 Pa, 1 kPa and 10 kPa were prepared in 21 mm glass-bottomed Petri dishes (35 mm, high; Ibidi) as described previously[11,23,24]. The gel premixes were prepared using predefined ratios of 60% phosphate buffered saline (PBS; Fisher Scientific), 40% (w/v) acrylamide (AA) solution and 2% bis-acrylamide (Bis-AA) solution (Fisher Scientific) as follows: ~0.1 kPa (5% AA and 0.04% Bis-AA), 1 kPa (7.5% AA and 0.06% Bis-AA), and 10 kPa (12% AA and 0.2% Bis-AA). The PAA gels were approximately 160 µm thick (40 µL of the premix spread across an 18 mm wide coverslip). One PAA gel of each stiffness was set aside for verification of stiffness using AFM.

Gels were functionalised with an overnight incubation of 100 µg/ml poly-D-lysine solution (PDL; MW 70,000-150,000), followed by 10 µg/ml laminin (from Engelbreth-Holm-Swarm murine sarcoma basement membrane) in culture medium (60% Leibovitz L15 medium, 1× PSF, pH 7.6-7.8) for 2 hours prior to culturing.

*Fabrication of 3D collagen gels*

3D collagen-based hydrogels were prepared as developed by Pillai, Mukherjee et al., 2024[5]. To create premixes for soft (G' = 40 Pa) and stiff (G' = 450 Pa) hydrogels, 6 mL of 25mM HEPES, 1× Leibovitz L15 medium, 1× PSF, pH 7.3 were diluted with 3 and 1 mL of distilled water, respectively. Throughout the protocol, the premixes, along with a stock of 10 mg/mL TeleCol-10 collagen (Advanced BioMatrix), were kept on ice to slow solidification. Stage 33/34 *Xenopus* brains were dissected out and left in culture medium (60% Leibovitz L15 medium, 1× PSF, pH 7.6-7.8) until sufficient brains were collected for one hydrogel. Collagen was then rapidly diluted in the appropriate premix to a volume of 1 mL, mixed through pipetting, and expelled into a glass-bottom Petri dish (35 mm, high; Ibidi). The dilution was as follows: ∼40 Pa: 1 mg/mL collagen (70% "soft" premix and 30% collagen) and ∼450 Pa: 3 mg/mL collagen (90% "stiff" premix and 10% collagen). The dissected brains were immediately pipetted into the gel droplet and the hydrogel was left to solidify for 2 hours at 20°C. Subsequently, enough culture medium was added to the dish to cover the solidified gel, and the sample was left at 20°C for another 22 hours. Samples were then fixed in modified Carnoy's fixative[6] for 2 hours and HCR RNA-FISH was carried out as described for whole-mount brains (Fig. 4a). During the dissection step of the HCR RNA-FISH protocol, the brains were dissected out of the 3D collagen gel instead of the embryo.

#### *Xenopus* retinal explant culture

Stage 35/36 embryos were washed briefly in 1× PSF (Lonza), anesthetised, and whole eye primordia were dissected out as previously described[11]. Whole eye primordia were immediately transferred to a 35 mm Petri dish coated with Sylgard-184 containing culture medium (60% Leibovitz L15 medium, 1× PSF, pH 7.6-7.8), where they were gently pipetted up and down 2-3 times. Once there were enough eye primordia for a single culture dish, eye primordia were dissected into thirds along the dorso-ventral axis to increase the precision in extracting axons expressing the highest and lowest EphB levels[25]. Eye primordial sections were then pipetted into the culture dish and pushed down lightly with dissection tools to increase adherence to the substrate. Dorsal and ventral sections were cultured in separate regions of the same dish to decrease the variability arising from human error between technical replicates. The process was repeated for all culture dishes. To allow for adhesion, dishes were left undisturbed for 30 min (glass) or 4 hours (hydrogels), after which they were transferred into 20°C for 24-25 hours.

#### Growth cone collapse assay

To allow for clustering of the ephrinB1 proteins, soluble recombinant mouse ephrinB1-Fc chimera protein (R&D Systems), or human IgG1 Fc peptide (Jackson Immunoresearch Labs) were incubated with a goat antibody against human IgG, Fc fragment (Jackson Immunoresearch Labs) as previously described[26,27]. Clustered ephrinB1-Fc and Fc only were bath applied to 24-hour retinal cultures for 30 min at a 1:1 ratio with the culture medium in the dish, to give a final concentration of 5 µg/mL. Cultures were then fixed in 2% PFA, 7.5% sucrose for 15 min, carefully washed three times with PBS, and stored at 4°C for up to 2 days before imaging. Samples were imaged on a Leica DMi8 inverted microscope with a

20×/0.4 phase contrast objective (ORCA-Flash4.0 V2 Digital CMOS camera Model C11440-22CU) or a Nikon ECLIPSE Ti microscope with a 20×/0.5 Plan Fluor Nikon phase contrast objective (Photometrics Prime BSI Scientific CMOS (sCMOS) camera). Analysis was carried out blinded, by manually counting the number of collapsed and healthy growth cones on acquired images (Fig. 1a). Generally, a growth cone is considered `collapsed' if it has two or fewer filopodia and no visible lamellipodia[28] (Fig. 1b-c). Only growth cones from individual axons or thin axon bundles were counted. A biological replicate refers to any number of embryos from the same *in vitro* fertilisation.

The data is additionally expressed as normalised percentage collapse, where the percentage of growth cone collapse in the ephrinB1 condition was normalised by the percentage collapse in the corresponding control (Fc) condition belonging to the same biological replicate. Specifically, the normalised percentage collapse is the difference between the percentage of growth cone collapse between the ephrinB1 and Fc conditions divided by the subtracted `baseline' collapse (100 minus the percentage of Fc collapse), multiplied by 100. This method of presenting the data removes variation of baseline level collapse due to differences in neuronal health between biological replicates.

**Analysis of fluorescence in whole-mount brains**

The following analysis method was used for the quantification of nuclear density and mRNA levels (HCR RNA-FISH). All images were processed in Fiji[29]. A maximum intensity z-projection was made, and the image was rotated so the A-P axis of the brain was aligned along the horizontal plane, with the anterior to the left. A brightfield image was used to extract only pixels belonging to the optic tectum from the fluorescence channel(s).

Specifically, the z-stack was first converted into 32-bit, a selection was created around the boundaries of the optic tectum. The region outside the selection was set to NaN so that only pixels within the optic tectum remained. A new selection was then made using the rectangle tool around the boundaries of the optic tectum, and the intensity profile across the horizontal axis was recorded for each fluorescence channel. The intensity profile was then normalised according to the minimum and maximum intensity per brain. This was performed for all biological replicates (*Xenopus* brains).

To plot intensity profiles for individual genes at different stages of innervation, a rolling mean of 20 µm was applied to each individual line profile, roughly equivalent to what a growth cone can sense *in vivo*[7]. The median mRNA level at each A-P position was then plotted, along with a ribbon denoting the 95% confidence interval. These intensity profiles were also used to calculate the position of the A-P peak mRNA expression.

The same ROI of the optic tectum was also used to calculate the absolute median mRNA levels between different embryos for HCR RNA-FISH (Fig. 3).

*Generating mean normalised expression maps of the optic tectum*

The A-P axis-aligned fluorescence images described above, along with the ROI of the optic tectum were also used to generate mean normalised maps of mRNA expression for each gene. This was done using custom-written a MATLAB script like that used for generating the mean normalised stiffness maps described previously (https://github.com/FranzeLab/AFM-data-analysis-and-processing/blob/master/SipkovaPaper2024/HCR_grid_tectum_analysis_FIRST.m). Briefly, the ROI generated in Fiji was used to select only the intensity values within the optic tectum of the aligned fluorescence images. These values were then converted into a coordinate system normalised from 0 to 1 to allow comparison between biological replicates with varying tectum sizes. Then, the same custom-written MATLAB script described previously (https://github.com/FranzeLab/AFM-data-analysis-and-processing/blob/master/SipkovaPaper2024/Avg_tectum_map_SECOND.m) was used assign each value to a grid coordinate, calculate the mean grey value associated with each grid point, and plot the dataset as a heatmap for each gene at each stage of innervation. To compare mRNA expression and stiffness in the optic tectum, the mean mRNA level and mean stiffness at each grid point were plotted against each other for each gene at each stage of innervation, and correlation analysis was performed.

**Statistical analysis**

All statistical tests and plotting were performed in R[30]. For each dataset, a Shapiro-Wilk and Levene's test were used to test for normality of the underlying distribution and equality of variances across the different conditions, respectively. Levene's test was chosen as it is less sensitive to departures from normality than other equivalent tests. Additionally, the normality and equality of variances were visually inspected using a quantile-quantile (Q-Q) plot and residual plot, respectively. For data which satisfied both assumptions, parametric tests were used, namely a Welch t-test or one-way ANOVA followed by a post-hoc Tukey test. Otherwise, the Mann Whitney U test or a Kruskal-Wallis chi-squared test followed by pairwise comparisons using a Mann Whitney U test with a Benjamini-Hochberg correction were used.

### Supplementary Figures

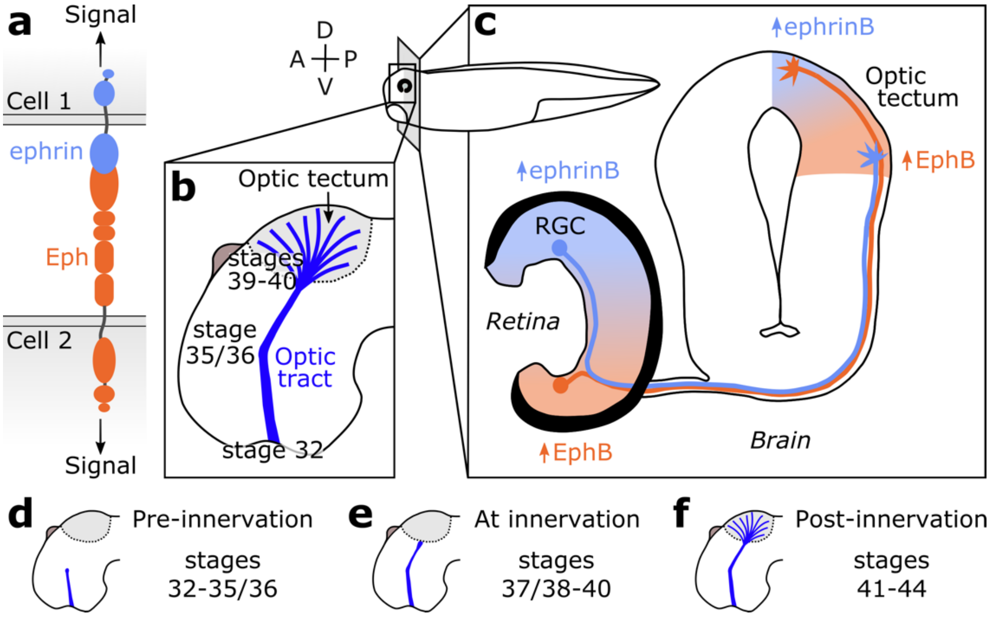


**Figure S1: Eph/ephrin signalling in the *Xenopus* retinotectal projection.** (**a**) General structure of the transmembrane proteins Eph (orange) and ephrin (light blue). When bound, signalling can be unidirectional or bidirectional. (**b**) Lateral view of a *Xenopus* brain at stage 48, when retinal ganglion cell (RGC) axons (blue) have innervated the optic tectum (light grey). Developmental stages of particular importance in the formation of the retinotectal projection are: 1) stage 32, when RGC axons cross the optic chiasm to grow along the lateral surface of the brain, 2) stage 35/36, when the optic tract makes a characteristic turn at the diencephalon-telencephalon boundary, 3) stages 39-40, when the first RGC axons reach the optic tectum, and 4) stage 48, when the initial ordered retinotectal map is established. Stages are according to Nieuwkoop and Faber (1994)[1]. (**c**) Schematic cross-section of the developing *Xenopus* retinotectal projection showing the counter-gradients of ephrinB and EphB expression across the retina and optic tectum. The dorso-ventral (D-V) axis of the retina is mapped onto the ventral-dorsal axis of the optic tectum. Image based on Erdogan et al., 2016[31]. Anterior (A), posterior (P), dorsal (D) and ventral (V). (**d-f**) Schematics of pre-, at and post- innervation stages of the optic tectum (light grey) by RGCs (blue). Throughout this paper, stages 32-35/36 are classified as pre-innervation, stages 37/38-40 as at innervation, and stages 41-44 as post-innervation.

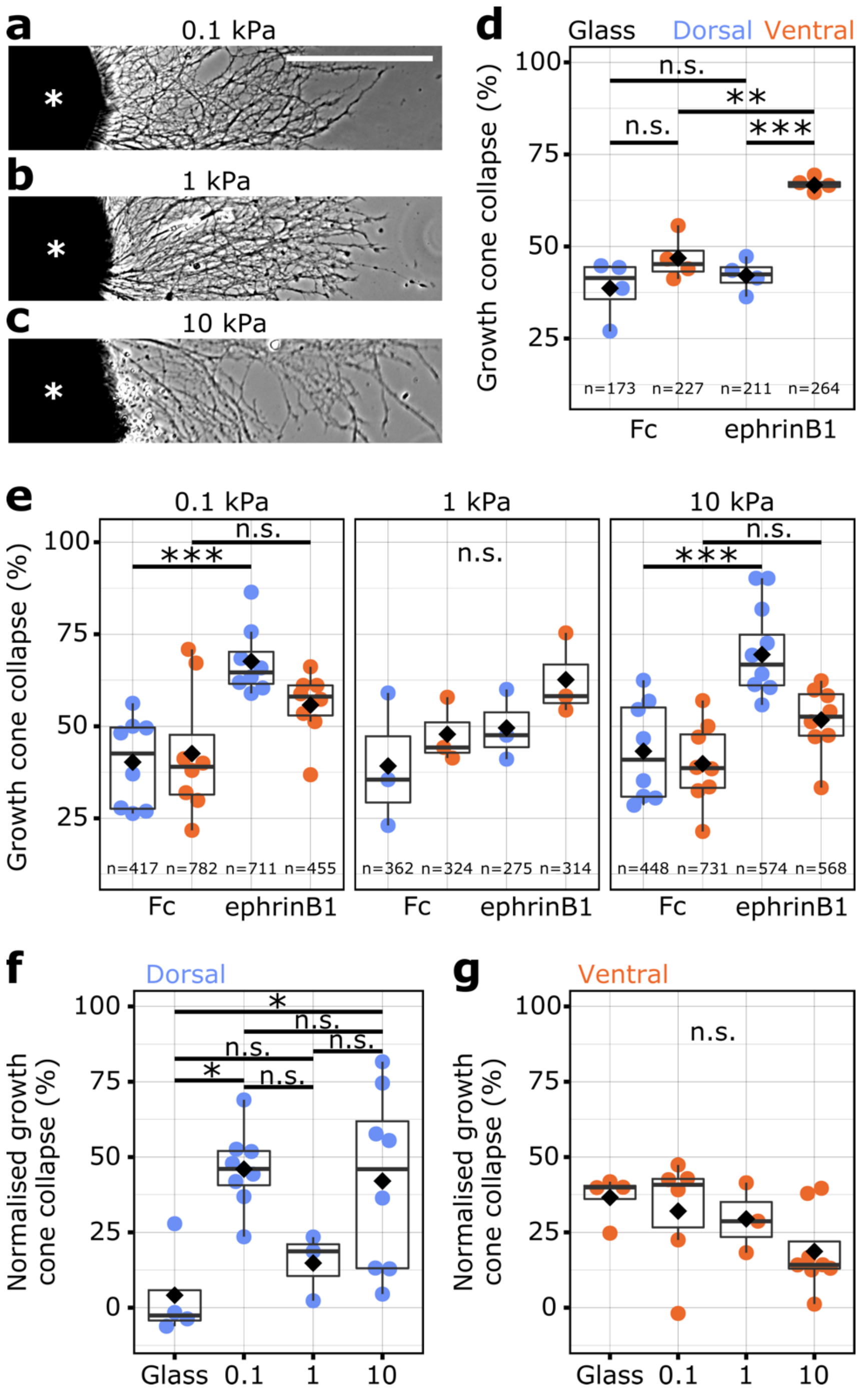


**Figure S2: Response of dorsal and ventral RGC growth cones to ephrinB on glass and compliant substrates.** (**a-c**) Representative images of whole eye primordial (marked with an asterisk) cultures on compliant substrates of 0.1, 1 and 10 kPa. Scale bar = 100 µm. The stiffer the substrates, the longer the axons. (**d-e**) The number of collapsed growth cones per condition expressed as a percentage of the total number of growth cones. (**d**) On glass, the response of dorsal and ventral RGCs to pre-clustered Fc domain fragments only (control)

was similar ($p$ = 0.235, one-way ANOVA followed by *post-hoc* Tukey test). Ventral RGCs responded more to ephrinB1 than dorsal RGCs ($p$ = 0.000292). When exposed to ephrinB1, ventral RGCs collapsed significantly more than in the control condition ($p$ = 0.00184), whereas dorsal RGCs did not ($p$ = 0.828). (**e**) In contrast, on substrates of 0.1 kPa and 10 kPa, dorsal RGCs responded more to ephrinB1 than the control condition ($p_{0.1\ kPa}$ = 0.000789, $p_{10\ kPa}$ = 0.000524, one-way ANOVA followed by *post-hoc* Tukey test), whereas ventral RGCs did not ($p_{0.1\ kPa}$ = 0.174, $p_{10\ kPa}$ = 0.191). There was no significant difference in response on 1 kPa substrates ($p$ = 0.227, F-value = 1.79, one-way ANOVA). Statistical analysis was conducted on each gel separately. (**f-g**) The percentage of collapse in response to ephrinB1 was normalised by the percentage of collapse in the corresponding control conditions. The normalised response of dorsal RGCs was significantly higher on 0.1 kPa and 10 kPa substrates than on glass ($p_{0.1\ kPa}$ = 0.0200, $p_{1\ kPa}$ = 0.909, $p_{10\ kPa}$ = 0.0381, one-way ANOVA followed by *post-hoc* Tukey test), whereas the normalised response of ventral RGCs did not differ between substrates ($p$ = 0.187, F-value = 1.79, one-way ANOVA). However, the response of dorsal RGCs did not differ between compliant substrates ($p_{0.1-1\ kPa}$ = 0.162, $p_{0.1-10\ kPa}$ = 0.981, $p_{1-10\ kPa}$ = 0.256). These data suggested that the observed lack of difference between the response of ventral and dorsal RGCs to ephrinB1 on soft substrates (Fig. 1e), which was different from the well-established response on glass, was due to an enhanced response of dorsal RGCs to ephrinB1 on soft substrates compared to glass. X-axis labels are expressed in kPa. In all boxplots, each point represents the mean collapse across a biological replicate (RGCs from embryos from the same *in vitro* fertilisation), n = number of growth cones. The black diamond denotes the mean. *p < 0.05, **p < 0.01, ***p < 0.001, n.s. = not significant.

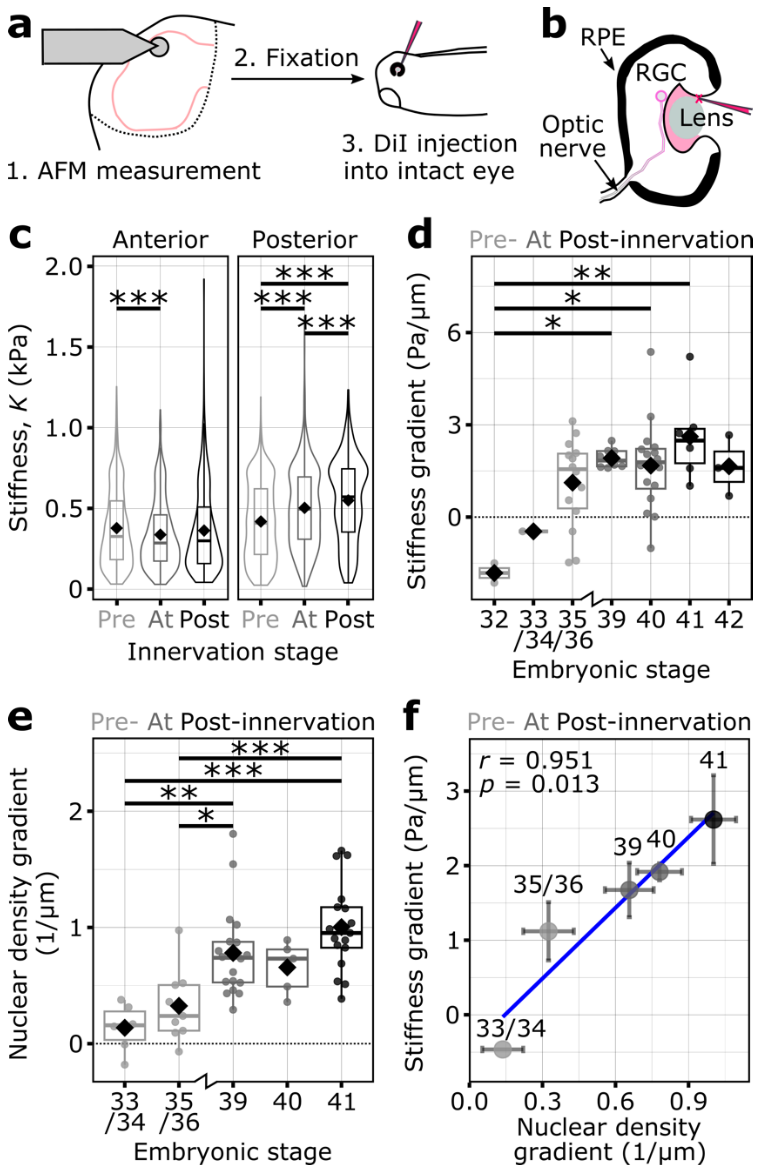


**Figure S3: Anterior-posterior stiffness and nuclear density gradients in the optic tectum correlate strongly across developmental stages during innervation by RGCs.** (**a**-**b**) Schematic of the AFM methodology, including DiI injection. (**c**) Comparison of stiffness values within the anterior or posterior half of the optic tectum between innervation stages. For clarity, due to the large number of measurements, a violin plot is used in conjunction with a boxplot to show the distribution of points. While the anterior region significantly softened between pre-innervation and at innervation stages ($p_{\mathrm{Pre-At}}$ = 0.00095, $p_{\mathrm{Pre-Post}}$ = 0.139, $p_{\mathrm{At-Post}}$ = 0.547, Kruskal-Wallis chi-squared test followed by *post hoc* Mann Whitney U test with a Benjamini-Hochberg correction), the posterior region stiffened between each innervation stage ($p_{Pre-At} < 10^{-15}$, $p_{Pre-Post} < 10^{-15}$, $p_{At-Post} = 0.00011$). The stiffness of the anterior and posterior optic tectum at pre-innervation were, respectively, A = 379 ± 7 Pa and P = 420 ± 7 Pa, at innervation: A = 338 ± 6 Pa and P = 504 ± 6 Pa, and post-innervation: A = 364 ± 14 Pa and P = 550 ± 12 Pa (mean stiffness ± standard error) (**d**) Stiffness gradients quantified across the A-P axis of stage 32-42 embryos. Negative values indicate a stiffer anterior region, while positive values indicate a stiffer posterior region. In general, younger embryos had lower stiffness gradient values ($p_{32-33/34} = 0.978$, $p_{32-35/36} = 0.0661$, $p_{32-39} = 0.0127$, $p_{32-40} = 0.0147$, $p_{32-41} = 0.00266$, $p_{32-42} = 0.0771$, $p_{33/34-35/36} = 0.901$, $p_{33/34-39} = 0.607$, $p_{33/34-40} = 0.690$, $p_{33/34-41} = 0.325$, $p_{33/34-42} = 0.797$, $p_{35/36-39} = 0.810$, $p_{35/36-40} = 0.905$, $p_{35/36-41} = 0.244$, $p_{35/36-42} = 0.995$, $p_{39-40} = 0.999$, $p_{39-41} = 0.953$, $p_{39-42} = 1.000$, $p_{40-41} = 0.738$, $p_{40-42} = 1.000$, $p_{41-42} = 0.940$, one-way ANOVA followed by *post-hoc* Tukey test). (**e**) Nuclear density gradients quantified across the A-P axis of stage 33/34-41 embryos. More positive values indicate higher pixel intensity values and thus more nuclei in the posterior region. In general, younger embryos had lower nuclear density gradient values ($p_{33/34-35/36} = 0.844$, $p_{33/34-39} = 0.00247$, $p_{33/34-40} = 0.117$, $p_{33/34-41} = 0.0000333$, $p_{35/36-39} = 0.0195$, $p_{35/36-40} = 0.442$, $p_{35/36-41} = 0.000191$, $p_{39-40} = 0.955$, $p_{39-41} = 0.343$, $p_{40-41} = 0.307$). For **c-e**, the mean at each stage is denoted by a black diamond. (**f**) The stiffness gradient and nuclear density values shown in **d** and **e** strongly correlated at each developmental stage (adjusted $R^2 = 0.87$, Pearson's correlation coefficient $r = 0.951$, $p = 0.013$, Pearson's product-moment correlation). The three different innervation stages (Fig. S1) are denoted by shades of grey. *p < 0.05, **p < 0.01, ***p < 0.001. Statistically insignificant relationships are not annotated.

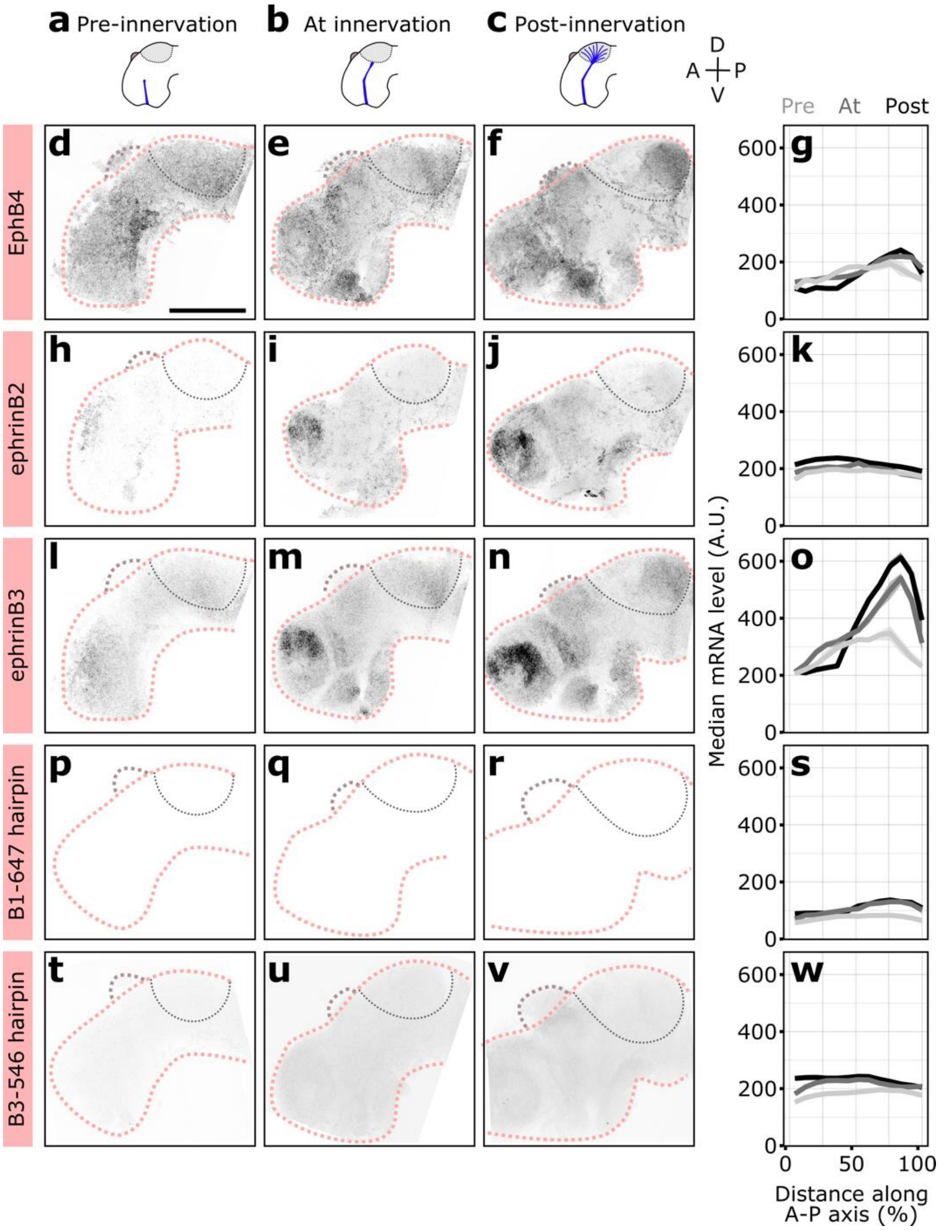


**Figure S4: EphB and ephrinB expression in the optic tectum during innervation by RGCs.** (**a-c**) Diagrams of the innervation stages of the optic tectum (light grey) by RGCs (blue) (Fig. S1). Anterior (A), posterior (P), dorsal (D) and ventral (V). (**left three columns**) Representative images of EphB4 (**d-f**), ephrinB2 (**h**-**j**) and ephrinB3 (**l**-**n**) mRNA expression in the optic tectum over time. Grey values scaled within each gene, image size scaled over all genes; scale bar = 200 µm. Representative images for the (**p-r**) B1-647 hairpin and (**t**-**v**) B3-546 hairpin used in conjunction with all EphB and ephrinB mRNAs, respectively. The negative control images are scaled as follows: the B1-647 hairpin to EphB2 and the B3-546 hairpin to ephrinB1 (genes shown in Fig. 3). (**g**, **k**, **o**, **s**, **w**) Median grey values across the normalised A-P axis of the optic tectum at the three stages of innervation for (**g, k, o**) probe and (**s, w**) negative control (hairpin only) conditions. EphB4 (**g**) and ephrinB3 (**o**) peak expression shifted to the posterior end of the tectum during innervation, whereas ephrinB2 (**k**) expression remained homogeneous throughout the optic tectum. The negative controls confirmed that non-specific binding of the B1-647 (**s**) and B3-546 (**w**) hairpins to the *Xenopus* brain was negligible at these stages. In each line profile, the median grey value of every 20 µm is indicated by a line and a ribbon denotes the 95% confidence interval. Biological replicates *n* for each gene at each stage are listed in Fig. S5. Embryos were obtained from three *in vitro* fertilisations.

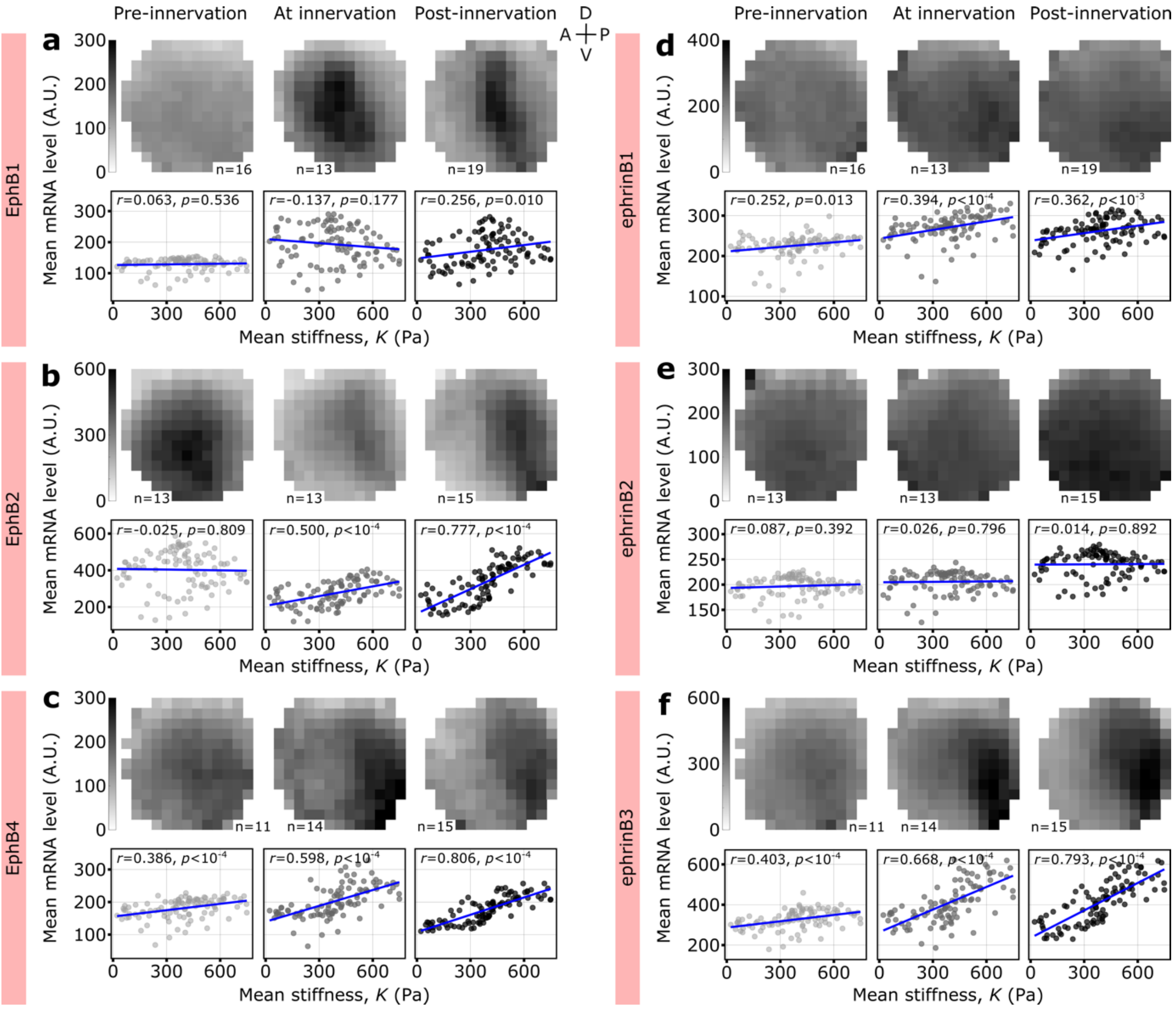


**Figure S5: Correlation analysis between EphB/ephrinB expression and tissue stiffness in the optic tectum.** (**Upper rows**) Mean normalised scaled maps of EphB1 (**a**), EphB2 (**b**), EphB4 (**c**), ephrinB1 (**d**), ephrinB2 (**e**) and ephrinB3 (**f**) mRNA expression in the *Xenopus* optic tectum across innervation stages (see Methods for quantification details; Fig. S1d-f). Anterior (A), posterior (P), dorsal (D) and ventral (V). Each pixel is the mean mRNA level (i.e. grey value) associated with that location in the tectum across *n* embryos. Embryos were obtained from three *in vitro* fertilisations (same data as in Fig. S4). (**Lower rows**) To directly compare mRNA expression and stiffness values in the optic tectum, the mean mRNA level at each pixel position was then plotted against the mean stiffness value at the same pixel position in the normalised optic tectum (Fig. 2g-i). This was done for each gene across all stages of innervation, and linear correlation analysis (blue line) was performed using Pearson's product-moment correlation (see Methods for quantification details). EphB2 (**b**), EphB4 (**c**) and ephrinB3 (**f**) expression correlated strongly with tissue stiffness (see also Table S1). In contrast, EphB1 (**a**) and ephrinB1 (**d**) showed only a weak positive correlation with tissue stiffness, and ephrinB2 (**e**) showed no correlation at all. Pearson's correlation coefficient, *r* and *p*-value are reported for each analysis.

### Supplementary Tables

| *Gene* | Innervation stage | | | | | | | | |
|---|---|---|---|---|---|---|---|---|---|
| | Pre-innervation | | | At innervation | | | Post-innervation | | |
| | $R^2$ | *r* | *p*-value | $R^2$ | *r* | *p*-value | $R^2$ | *r* | *p*-value |
| ***EphB1*** | -0.006366 | 0.06332 | 0.5357 | 0.008616 | -0.1368 | 0.1767 | 0.05596 | 0.2559 | 0.01017 |
| ***EphB2*** | -0.009696 | -0.02464 | 0.8087 | 0.242 | 0.4997 | 1.396e-07 | 0.5992 | 0.7767 | < 2.2e-16 |
| ***EphB4*** | 0.1402 | 0.3858 | 7.368e-05 | 0.3509 | 0.5978 | 5.126e-11 | 0.6465 | 0.8063 | < 2.2e-16 |
| ***ephrinB1*** | 0.05353 | 0.2516 | 0.01246 | 0.1464 | 0.3938 | 5.506e-05 | 0.1222 | 0.3620 | 0.0002146 |
| ***ephrinB2*** | -0.002677 | 0.0869 | 0.3923 | -0.009607 | 0.02637 | 0.7956 | -0.01001 | 0.01375 | 0.892 |
| ***ephrinB3*** | 0.154 | 0.4032 | 3.189e-05 | 0.4406 | 0.6680 | 3.146e-14 | 0.6242 | 0.7925 | < 2.2e-16 |

**Table S1: Correlation analysis between EphB and ephrinB expression and tissue stiffness in the optic tectum during innervation by retinal ganglion cells.** Adjusted $R^2$, Pearson's correlation coefficient, *r,* and p-value from the linear regression and Pearson's product-moment correlation analyses conducted on data in Fig. S5.
